# Optimisation of surfactin yield in *Bacillus* using active learning and high-throughput mass spectrometry

**DOI:** 10.1101/2024.01.24.576661

**Authors:** Ricardo Valencia Albornoz, Diego Oyarzún, Karl Burgess

## Abstract

Integration of machine learning and high throughput measurements are essential to drive the next generation of the design-build-test-learn (DBTL) cycle in synthetic biology. Here, we report the use of active learning in combination with metabolomics for optimising production of surfactin, a complex lipopeptide resulting from a non-ribosomal assembly pathway. We designed a media optimisation algorithm that iteratively learns the yield landscape and steers the media composition toward maximal production. The algorithm led to a 160% yield increase after three DBTL runs as compared to an M9 baseline. Metabolomics data helped to elucidate the underpinning biochemistry for yield improvement and revealed Pareto-like trade-offs in production of other lipopeptides from related pathways. We found positive associations between organic acids and surfactin, suggesting a key role of central carbon metabolism, as well as system-wide anisotropies in how metabolism reacts to shifts in carbon and nitrogen levels. Our framework offers a novel data-driven approach to improve yield of biological products with complex synthesis pathways that are not amenable to traditional yield optimisation strategies.

**Graphical abstract:** 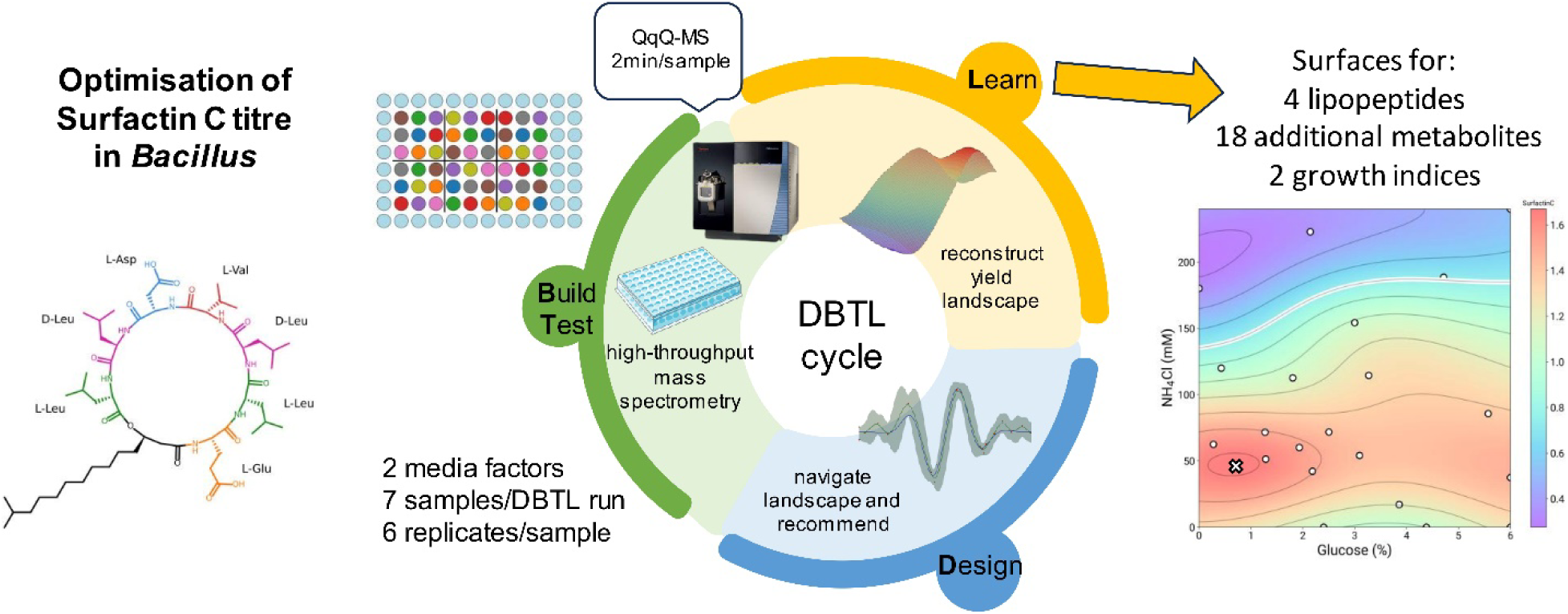

## Introduction

Surfactants are compounds that lower the surface tension between two liquids or a liquid and gas. Their steady demand in industrial and domestic applications has motivated searching for alternatives to traditional petroleum-derived compounds, so as to reduce long-term environmental impacts [1]. Recent years have witnessed a growing number of approaches for bio-surfactant production that are environmentally friendly, safe for human health, and biodegradable [2-4].

Surfactin is a promising bio-surfactant suitable for high temperature conditions required in industrial applications [5-10]. It is a lipopeptide produced by several strains of the genus *Bacillus* [11] with a wide range of applications in industry, agriculture, and medicine, such as emulsifiers, dispersants, and biocontrol agents [12-15]. As an endogenous metabolite, surfactin facilitates cell motility and colonization [16-18]. When interacting with other bacteria, it functions as a spatially distributed antibiotic that disrupts the membrane of nearby cells, a phenomenon that can also promote the emergence of surfactant-resistant communities [19-22].

Surfactin biosynthesis is carried out by a non-ribosomal peptide synthetase (NRPS) [8, 23, 24]. Due to its combinatorial assembly mechanism, surfactin production depends on multiple pathways that supply the molecular precursors for assembly in the cytoplasm. Such precursors include branched-chain fatty acids and amino acids [25, 26]. These multiple production routes confer substantial flexibility to the biosynthetic processes and translate into the production of several variants of surfactin [12], especially if a precursor undergoes modifications or if there are errors in the amino acid assembly.

Key challenges for surfactin production are the high variability and low titres observed in liquid medium, which introduce substantial barriers to scale up production [25,26]. Current strategies to increase production include genetic engineering and optimisation of fermentation conditions [25, 26]. For example, overexpression of the efflux pump *swrC* (synonym: *yerP*) and other pumps has been reported, deriving into an 0.5-1.5-fold increase of surfactin production [25]. Additionally, overexpression of the *P*_srf_ promoter or its substitution with a high expression promoter resulted in titres between 0.04-1.5 g/L [25], while efforts in metabolic engineering have focussed on rewiring *Bacillus* metabolism to generate surfactin hyperproducers [27]. Several other studies have employed design-of-experiment approaches to maximise surfactin titres by altering the concentrations of the medium components [28, 29]. Components of the media described in the publications include laboratory-grade chemicals and more complex substrates from industrial waste [30-34]. Although various studies have identified pathways that contribute to surfactin synthesis in *B. subtilis* [35] and *B. velezensis* [36], yield improvements in surfactin production have remained elusive thus far.

Here, we developed a machine learning pipeline to increase surfactin yield via iterative improvements to the media composition in a design-built-test-learn (DBTL) loop. Our approach utilizes mass spectrometry to quantify surfactin as well as a suite of background metabolites that provide insights on the metabolic processes that drive the yield increase. We constructed a data-efficient active learning loop whereby *Bacillus* cultures are grown in different media compositions that are guided by a machine learning predictor of the yield landscape. Active learning is a machine learning paradigm whereby models are trained on iterative batches of measurements; it is particularly useful in applications where comprehensive coverage of the input space is prohibitively expensive, and thus ideally suited for DBTL loops that involve mass spectrometry readouts. Through iterative DBTL rounds of measurement, model training and querying, we were able to improve surfactin yield by 160% as compared to a baseline M9 media.

We employ a Bayesian optimisation routine [37-40], a common tool for black-box optimisation that is commonly employed for tuning deep learning models [41, 42]. Recently, synthetic biologists have begun to explore its application in various use cases; early examples include synthetic gene design [43], as well as automated optimisation of metabolic engineering tasks [44-47], media optimization in cell-free systems [48], an *in silico* design of genetic control circuits [49]. Various bespoke approaches have been developed for media optimisation in mammalian cell bioproduction [50-52], and recently several software packages have been developed for the optimisation of gene circuits and metabolic pathways [53]. Our work is a novel application of active learning in combination with metabolomic readouts and thus offers substantial opportunities for data- and cost-efficient optimisation of complex bioprocesses across scales.

## Results

### Active learning of the surfactin C yield landscape

Our strategy to improve titre focussed on optimisation of culture conditions to enhance production of Surfactin C in the spent media from *Bacillus subtilis* DSM 3256 using an iterative active learning loop (Figure 1A). The medium composition variables, specifically the carbon concentration sourced from glucose and nitrogen concentration derived from ammonium chloride, were adjusted against the background of a baseline M9 medium. While temperature and agitation are pivotal to surfactin production [28], in this study the were kept constant to simplify the experimental design. The active learning cycle tested seven different carbon and nitrogen concentration combinations for Surfactin C production in each iteration. Each iteration can be seen as a different run of the Design, Build, Test, and Learn cycle in synthetic biology (Figure 1A).

**Figure 1.**
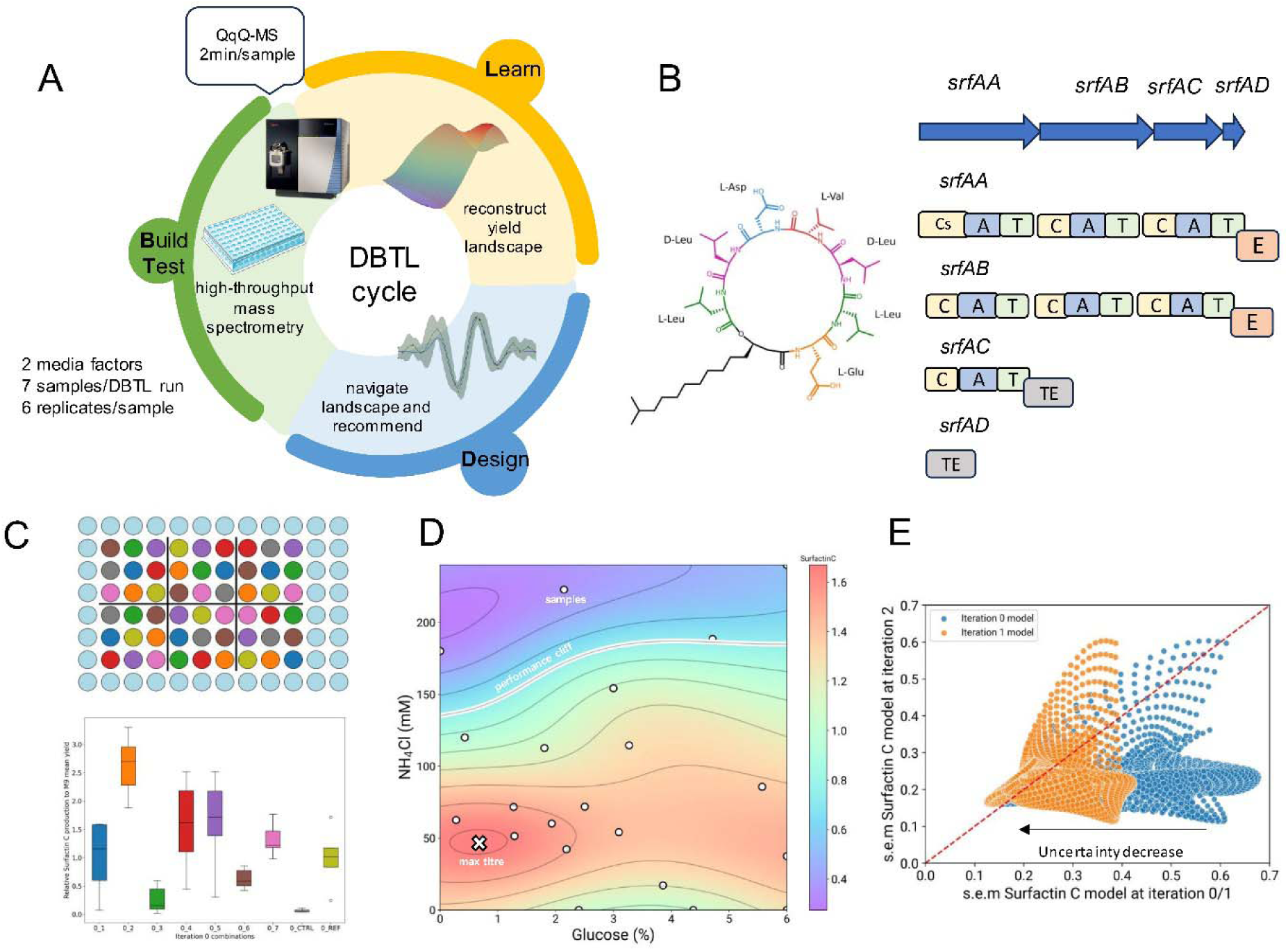
Active learning strategy for media optimisation. **A)** Active learning loop embedded in the Design, Build, Test, Learn (DBTL) cycle. The Build and Test stages encompass the cultivation of bacteria in microplates using machine learning-suggested combinations, followed by metabolite measurements via mass spectrometry. In the Learn stage, a Gaussian process regressor serves as a surrogate model, forecasting the Surfactin C titre landscape together with its prediction uncertainty. The Design phase, meanwhile, focuses on proposing new combinations, aided by the acquisition function. **B)** The structure of Surfactin C is displayed, emphasising the ring amino acids. The sequence for these amino acids is L-Glu1-L-Leu2-D-Leu3-L-Val4-L-Asp5-D-Leu6-L-Leu7. A simplified surfactin biosynthetic cluster diagram presents the four genes within the cluster and their corresponding domains: C for condensation; A for adenylation; T for thiolation or peptidyl carrier protein (PCP); E for epimerase; and TE for thioesterase. The initial condensation domain Cs in *srfAA* facilitates the integration of the fatty acid at the synthesis commencement. **C)** This section shows the layout for the microplate experiment during the initial iteration of active learning. Combination treatments, alongside control and M9 reference treatments, underwent block randomisation, resulting in six blocks. These are associated with biological replicates. A boxplot displays the Surfactin C titres achieved in the initial iteration. **D)** The final Surfactin C landscape following all three iterations is shown. The combination predicted to yield the maximum titre is marked with a cross. A zone exhibiting reduced titre (less than 0.5 in relation to the M9 titre, known as the performance cliff) is delineated with a white line. **E)** A dense grid of carbon/nitrogen combinations was employed to estimate uncertainty levels, expressed as the standard error of the mean, for models updated after iterations 0, 1 and 2. Models are compared pairwise, showcasing the uncertainty for identical grid points when comparing the model from iteration 2 against those from iterations 0 and 1.

In this study, the Build and Test stages involve cultivating the bacteria in microplates with varying combinations of glucose and ammonium chloride and then measuring the resulting metabolites using mass spectrometry. Among the 22 metabolites measured, the key product Surfactin C (Figure 1B) and three other lipopeptides (Surfactin B, Surfactin D, and Iturin A) were assessed using a flow injection-mass spectrometry method between iterations (Figure S1). Although a few colourimetric methods have been developed for surfactin and general lipopeptide quantification [54-58], we opted for mass spectrometry to provide a direct platform for the optimisation and it can therefore be generalised to compounds where a colourimetric/biosensor methods do not exist. This method was developed specifically for the experiment, given its ability to measure a single sample within 2 minutes and a full 96-well microplate in approximately 3 hours, thus streamlining the process.

Microplate experiments were randomized using a custom script in Python that can be used that can be generalised to other systems (see Data Availability). Optical density (OD) was measured (Figure S3, Figure S6) and Surfactin C titre results for iteration 0 are depicted in Figure 1C. We opted to limit the number of combinations per plate in favour of a larger number of replicates of each point (n=6) so as to accurately capture and incorporate biological variation into the machine learning model. The initial media compositions in Figure 1C were derived from a Latin hypercube design.

In the Learn phase, we employed a Gaussian process regressor as a surrogate model that predicts surfactin C titre and its uncertainty landscape in a 2-dimensional input space of media compositions. For the Design stage, we queried the model for new media compositions determined from a suitably chosen acquisition function that balances exploitation, i.e. select high-titre locations in the landscape, and exploration, i.e. cover areas of the design space where model predictions remain uncertain. Details of the active learning loop can be found in the Methods.

As shown in Figure 1D, we obtained a 160% titre improvement after three DBTL iterations improvement relative to the baseline titre in M9 medium. The optimal titre was reached at 0.8% glucose and 50 mM NH_4_Cl. Previously reported values for optimal surfactin production, including modifications to the Cooper [59] and Landy media regarding carbon and nitrogen concentrations, correspond to 0.8% glucose and 100 mM NH_4_Cl [60], in agreement with the obtained maximum for carbon concentration. When updating the models after each iteration (Figure S4), the average uncertainty in the predictions decreases from 0.45 to 0.3 (Figure 1E), indicating that the model gains more information about the system across iterations.

Thanks to our metabolomics measurement, we were also able to examine the yield of other surfactin variants (Surfactin B and Surfactin D), as well as the lipopeptide Iturin A which is assembled by a different biosynthetic cluster. We found that their production profiles are similar to that of Surfactin C (Figure S1), which is consistent with previous reports on co-production of fengycin and surfactin [61] and has also been observed in other categories of biosynthetic gene clusters [62].

To further examine the cellular constraints that affect surfactin production, we investigated trade-offs between lipopeptide production and maximal growth *in silico* (Figure 2). Using the final machine learning models, trained on three DBTL runs, we simulated hundreds of media compositions and constructed Pareto-like constraints between lipopeptide titre and the maximum optical density (OD), which was also simulated from a respective model. The results show no evident trade-off for Surfactin C and Surfactin B in relation to growth (Figure 2A), which suggests that Bacillus can achieve high titre with minimal sacrifice in biomass. However, we observed a more pronounced trade-off for Surfactin D and Iturin A (Figure 2B), where an ∼5% increase in titre is associated with ∼10% reduction in growth. This suggests a diversion of metabolic resources away from growth toward lipopeptide production.

**Figure 2.**
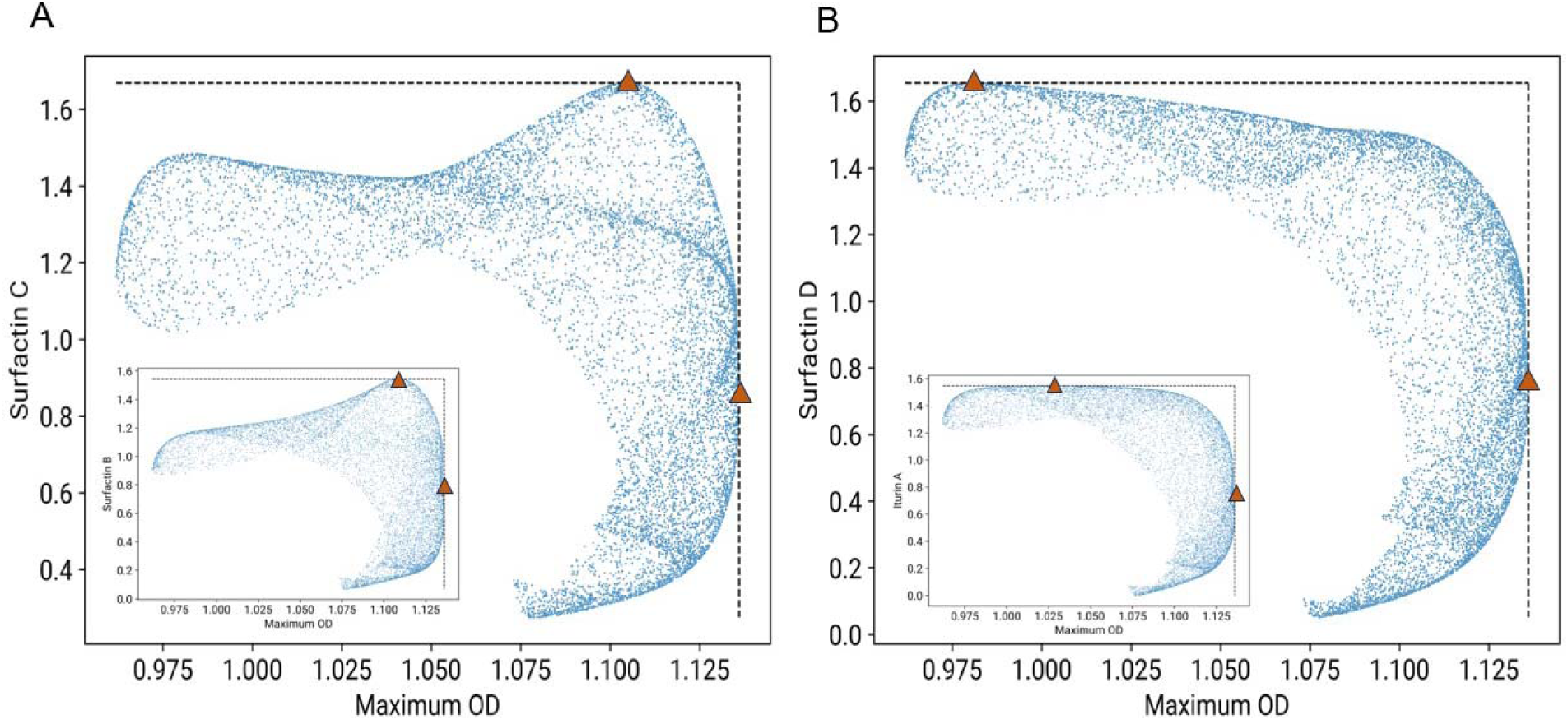
Trade-offs between lipopeptide yield and growth. **A)** For Surfactin C and D, growth can be minimally sacrificed to obtain a maximum of production, therefore, no trade-off is observed **B)** For Iturin A and Surfactin D, the Pareto front indicates an evident trade-off between lipopeptide production and bacterial growth. The maximum production of Iturin A and Surfactin D occurs at lower growth rates. In both cases the sample points were computed from simulations of lipopeptide production and growth with the final Gaussian process regressor model using a randomized sampling of the input space. Limits of the Pareto fronts are shown using red triangles.

### Impact of lipopeptide production on carbon-related metabolites

In addition to the lipopeptides, metabolites associated with carbon metabolism and the tricarboxylic acid cycle (TCA) can be measured using the flow injection method (Figure S2). Although these are not included in the medium, we were able to detect them in the spent medium because of export processes and release of cytoplasmic content from membrane disruption caused primarily by surfactin itself.

### The metabolic features of the loop samples show a relationship with surfactin production, and highlight the diversity of the samples

Using data from the loop samples alone, we performed a correlation analysis amongst the 4 lipopeptides, 18 other metabolites, and 2 other signals derived from growth data (maximum OD and final OD). A hierarchical clustering dendrogram highlights two primary clusters of metabolites, further divided into four correlated subgroups (Figure 3A). The first (6) and second groups (7) of the primary cluster contain a combination of amino acids and additional metabolites from the glycolysis/glucogenesis pathway. In contrast, the third (7) and fourth groups (5) consist of lipopeptides (4), and organic acids related to carbon metabolism (6), respectively. Both Canonical Correlation Analysis (CCA) and a PERMANOVA tests validated the distinct nature of these groups (CCA: p-value ∼1e-5 across all dimensions; PERMANOVA: p-value 1e-4). The organic acids in the fourth group show a positive correlation with lipopeptide production and are components or affiliates of the tricarboxylic acid cycle, suggesting heightened activity in this pathway.

**Figure 3.**
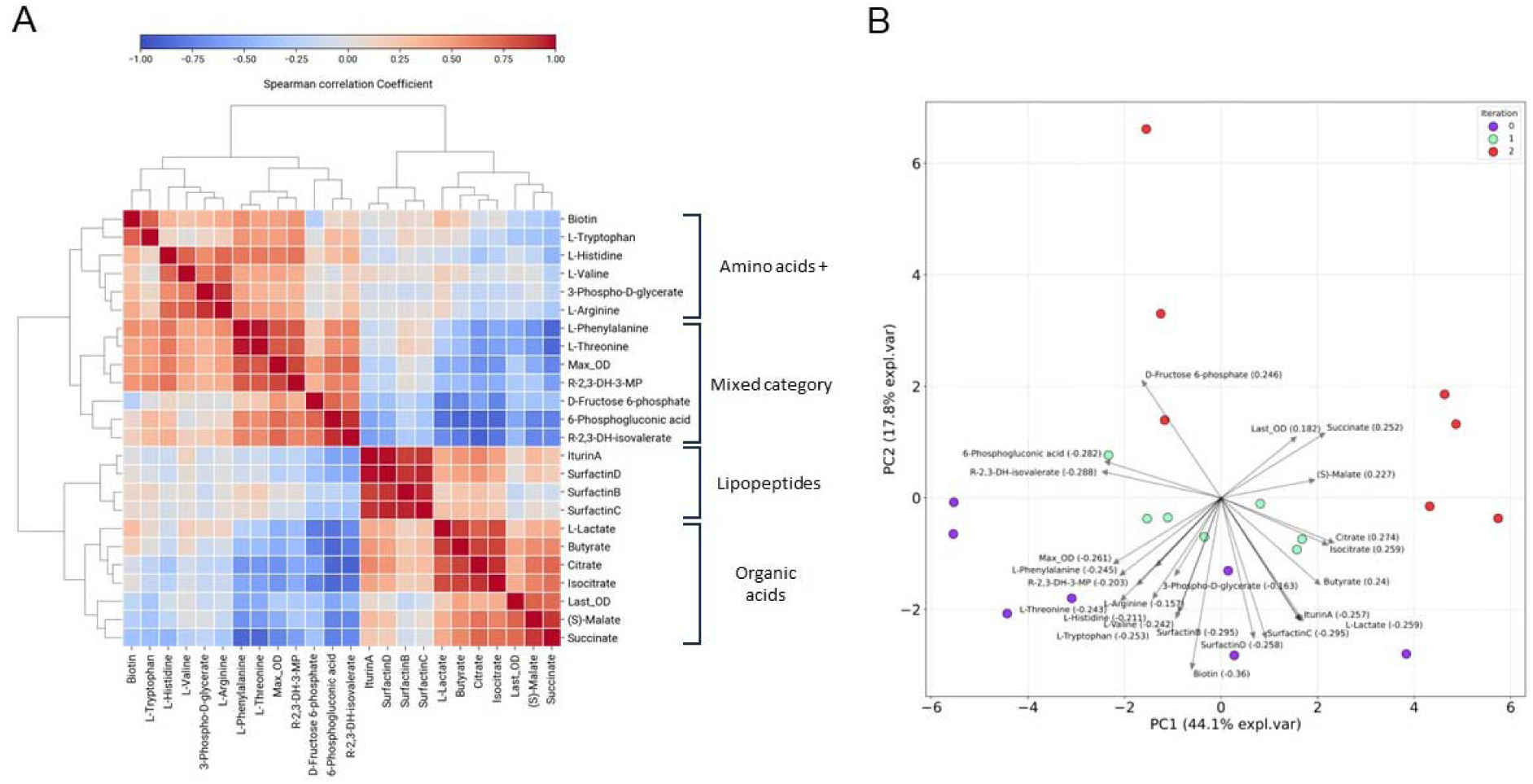
Analysis of background metabolic changes using the mass spectrometry data. **A)** Spearman correlation between the measured metabolites/growth across loop samples. The rows and columns are ordered by hierarchical clustering, shown as a dendrogram. Red colours indicate high positive correlation, while bluer colours indicate high negative correlation. Two primary clusters were identified from the dendrogram, which can be subdivided into 4 groups. Each of the groups is labelled within general biochemical categories. **B)** PCA biplot of the active learning loop samples. Iteration samples are depicted as points with distinct colours; blue for iteration 0, orange for iteration 1, and green for iteration 2. The arrows in the biplot represent the loading scores of each metabolite, indicating their influence on the principal components. This information is also presented in the scores accompanying the metabolite labels. To facilitate reading in the biplot, the metabolite names are abbreviated when necessary.

We performed a Principal Component Analysis (PCA) (Figure 3B) to explore the metabolic diversity of samples across different iterations, and additional relationships between metabolite measurements. The first and second components account for 44.1% and 17.8% of the explained variance, respectively. Notably, the lipopeptides and TCA cycle-associated organic acids account for the variance in the positive values of the first PCA component. In contrast, the metabolites from the primary cluster in Figure 3A show the opposite trend. The associated loading scores are similar in magnitude, indicating that no individual metabolite dominates the variance contribution after the reduction. Moreover, the sparsity of the points in the PCA space suggests that samples are metabolically diverse. To visualise the conclusions from the previous correlation/PCA analysis about how TCA metabolites have a similar landscape, and why the opposite trend is found in other metabolite categories, we embedded the metabolite surfaces into a simplified pathway diagram for anabolism in *Bacillus subtilis* (Figure S5) [63]. These are the predictions of the machine learning model for the abundance of each metabolite across the carbon and nitrogen concentration ranges. We found that L-arginine, which is linked with 2-oxoglutarate and is in the same pathway with glutamate synthesis, an amino acid in surfactin, possesses high abundancy with higher carbon-nitrogen concentration, suggesting higher activity in the glutamine synthetase-glutamate synthase (GOGAT) cycle and overflowing of this compound to the medium [64, 65].

### Sensitivity of metabolite production to changes in media composition

To quantify how metabolism reacts to simultaneous changes in carbon and nitrogen, we performed a novel directional analysis, whereby the direction in which carbon and nitrogen changes might have specific effect in the metabolic enzymes and reactions that can be measured with reference to the production surfaces generated. We term this concept as “metabolic anisotropy”, *i.e.*, non-uniformity in different directions or specifically in experimental terms, in simultaneous changes in medium composition. The approach consists of starting at the observed maximum of surfactin titre and tracing orthogonal trajectories to the level curves in different angular directions, until they reach a specified radius. This allows us to calculate the gradient of metabolite/growth/lipopeptide levels on angles from 0° to 360° respective to the Surfactin C maximum (Figure 4). For example, 0° corresponds to only increasing glucose, while 90° corresponds to only increasing NH_4_Cl concentration. Taking a radius of 0.6% glucose and 24 mM of nitrogen around the maximum carbon/nitrogen models, we used the trained models to calculate the gradient along the depicted blue circle (Figure 4A). A radar chart shows that several metabolites show near symmetric profiles, i.e., the rate of change of its abundance in every direction is similar, while others possess unusual long gradients for certain directions (Figure 6B). After filtering the gradient profiles that possess overall symmetry across every angle, we found 6 metabolites which present an asymmetric gradient profile, i.e., when stepping down of the optimum carbon/nitrogen concentration and moving towards another combination in a specific direction, the decrease/increase on that metabolite production is significantly different to when choosing another direction. Specifically, L-Phenylalanine, 6-Phosphogluconic acid and R-2,3-Dihidroxy-isovalerate exhibits a higher gradient (abundance change) at the 135° direction, corresponding to decreasing glucose concentration and increasing nitrogen concentration (Figure 6C). On the other hand, Succinate, (S)-Malate, and L-Arginine show higher gradients on the right half of the angle plane, moving towards increasing glucose concentration (Figure 6C). L-arginine, as we have shown before, has a production profile strongly associated with increasing carbon/nitrogen, pointing to a rapid response from nitrogen metabolism. The observed anisotropy on abundance changes for specific metabolites in the media could be related to multiple factors, including rapid enzymatic action, transient metabolic fluxes, overexpression of exporting systems, among others, and has not been thoroughly studied before on this experimental context, as far it is known. However, further experimentation is needed to confirm these profiles.

**Figure 4.**
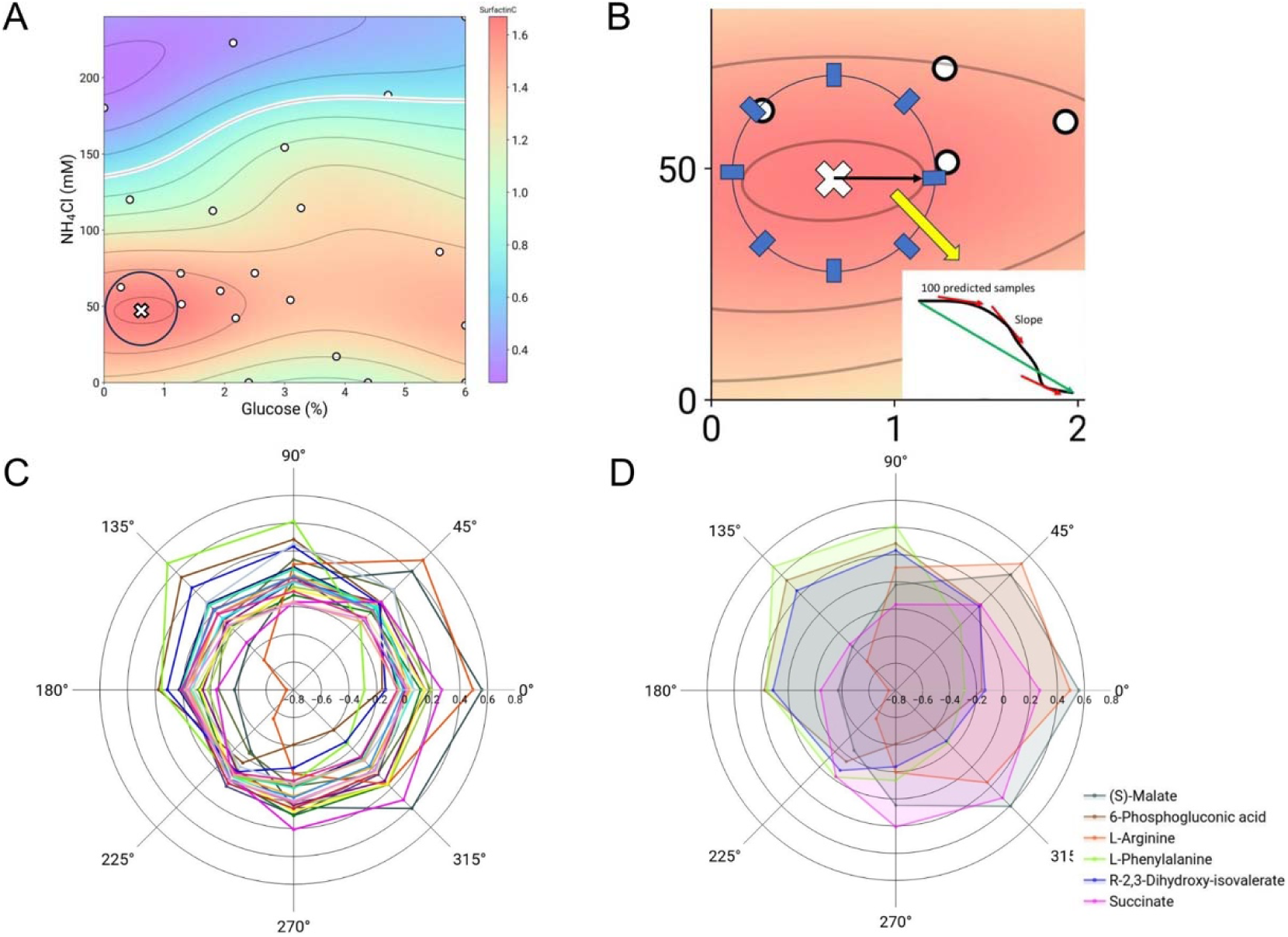
Directional analysis reveals metabolites with high sensitivity to changes in carbon and nitrogen composition nearby the Surfactin C maximum. . **A)** Radius around the maximum titre in the Surfactin C surface to calculate gradients in each metabolite production surface **B)** The gradients are calculated by simulating 100 samples in a path between the maximum titre point and a specific direction in the defined radius. Then the average gradient is obtained by averaging the (approximated) slopes for these samples. **C)** Profile of average gradient for different directions depicted as a radar chart. Each colour corresponds to a specific metabolite. The production surface for each metabolite was used to make the calculations. **D)** For specific metabolites, the profile is highly asymmetric, suggesting flux redistribution or enzyme kinetic changes when changing media composition simultaneously in a specific direction. The color legend is associated to each metabolite: (S)-Malate, 6-Phosphogluconic acid, L-Arginine, L-Phenylalanine R-2,3-Dihidroxy-isovalerate Succinate.

## Discussion

Large scale surfactin production in a liquid culture remains a challenge in bioprocessing [8], since higher titres are associated with lower growth. Therefore, we identified a need for high-throughput analysis to optimize surfactin production and overcome the trade-off between cell growth and biosynthesis. However, this should be accompanied by measurement technology that can keep with the pace of the experimental pipeline, as we exemplified with flow injection mass spectrometry and advanced statistical methods that enable human-readable data inspection. The methods and recommendations included in the paper have the potential to accelerate the development of DBTL-active learning platforms. The detailed picture of the rich metabolic data embedded in surfactin-associated pathways agrees with the literature [25, 26] and provides a powerful guide for further applications in metabolic engineering. Titre variation is a fundamental factor to consider when performing a Bayesian optimisation loop [41, 66]. Sources of noise include the stochasticity of the underlying biological process, errors introduced by liquid handling, instrumental variability, among others. Fundamentally, one can establish a trade-off where the total number of samples is reduced and the number of replicates per batch is increased, to gain confidence in the model’s predictions and thus achieve sample-efficient optimisation and reliable production data that can be used in downstream analysis. In addition, the selection of an appropriate model and acquisition function to account for this noise is paramount. Thus, our approach with 6 biological replicates, but less combinations per plate than reported active learning experiments, and the use of a specific acquisition function can greatly help the understanding of the system through the iterations.

To our knowledge, our study employs a novel combination of metabolomics and active learning for the optimisation of a bioprocess. Using mass spectrometry (MS) as the detection platform is a general approach, since allows the optimisation of titres for compounds which a fluorometric/colorimetric method or a biosensor are not available. In this context, MS can be complemented with high-throughput multi-omics measurements. This idea has been intensively developed over the last years and they are an effective complement in the development of a bioprocess optimisation strategy that is at the same time metabolically informative [67]. These measurements are already contributing to synthetic biology and metabolic engineering [68], but more effort is needed to integrate this data in the algorithms and machine learning models used by active learning loops.

## Methods

### Strains and media

*Bacillus subtilis* DSM 3256, a surfactin-producing *Bacillus* strain, was acquired from the Deutsche Sammlung von Mikroorganismen und Zellkulturen (DSMZ) repository. M9 medium was prepared using the following recipe: Glucose: 0.4% (carbon source), NH_4_Cl: 18.7 mM (nitrogen source), NaCl: 8.5 mM, MgSO_4_: 2 mM, CaCl_2_: 0.1mM, Na_2_HPO_4_: 42.2 mM, KH_2_PO_4_: 22 mM [69]. For the optimization experiments, glucose and ammonium chloride were excluded from this recipe, giving a basal M9 salts/buffer solution to add the carbon/nitrogen sources later. Components of the M9 medium were purchased from Sigma Aldrich. The microplate experiments were performed using ultra-low attachment surface flat bottom 96-well microplates (Corning).

### Microplate cultures

*Bacillus subtilis* DSM 3256 frozen stock was revived by culturing on an LB agar plate at 37°C overnight. From this plate, individual colonies were inoculated into six precultures consisting of 5ml of M9 medium. The cells were then grown in a shaking incubator at 37°C and 180 rpm.

Conditions corresponding to the carbon-nitrogen concentrations in the medium, as well as controls, were block-randomised across a 96-well microplate, considering six replicates (blocks). These were manually pipetted into place. The layout for block randomisation was obtained from an R script utilising the agricolae package v.1.3-7 [70]. In each well, 100µl of 2X M9 salts (M9 medium without a carbon or nitrogen source), 80µl of the glucose/ammonium mix, and 20µl of the preculture were added. This was adjusted to achieve an initial optical density at 600nm (OD600) of 0.1 in each well. The microplate culture was conducted using a Tecan Infinite M200 PRO plate reader, recording an OD600 measurement every 10 minutes for 36 hours at 37°C. The agitation was set to a maximum of 10mm. After culturing, the microplate was centrifuged for 20 minutes at 4000 rpm and 4°C to separate the cell pellet from the supernatant. 80µl of supernatant was taken from each well and stored in a -80°C freezer until mass spectrometry quantitation.

### Surfactin and metabolite quantification

We utilised flow injection of spent media with a ThermoFisher Dionex Ultimate 3000 autosampler. The sample volume was set at 1 μl. The mobile phase comprised a 1:1 ratio of acetonitrile to water with 0.1% formic acid, and the flow rate was 200 μl/min. The sample acquisition time spanned 1 minute, using selected reaction monitoring (SRM) on a ThermoFisher TSQ Quantiva triple quadrupole (QqQ) mass spectrometer (MS). The MS parameters, as well as the precursor/product masses table, can be found in Supplementary Tables 1 and 2, respectively.

Peak extraction was accomplished using the rawrr package v1.10.1 in R [71]. Subsequently, metabolites that needed it underwent baseline correction using the asymmetric least squares algorithm from the Python pybaselines package v1.0.0 [72] set to default parameters. The corrected peaks were integrated over the 1-minute run using the trapezoid rule function from the Numpy package via a custom Python script. The integrated intensity data was then compiled into a table.

Outliers were identified and removed based on the interquartile range (IQR), keeping only values within the range of median - 1.5IQR to median + 1.5IQR. These values were then normalised by dividing them by the average M9 titres observed for each batch. A table comparing the relative surfactin C titre to the observed M9 titre for every condition or combination was constructed for the subsequent active learning prediction step. Similar tables were generated for additional metabolites.

### Active learning loop

From the minimum and maximum concentrations of glucose and ammonium chloride that were selected for testing, a 2D design space was defined. Seven initial conditions were obtained from a Latin hypercube design (LHD). The centred LHD was implemented in Python using the pyDOE2 package [73]. After getting the relative surfactin C titre to the observed M9 titre from the MS quantification, these values were fitted using a heteroskedastic Gaussian process regression (GPR) model [74, 75], where the corresponding glucose/ammonium concentrations are inputted as variables/features and the titre is the observed output. From the GPR predictions, the q-Noisy expected improvement (q-NEI) acquisition function [66,75] is calculated for the design space, and optimising this function retrieve seven combinations to be tested on the next iteration of the loop. The model and the acquisition function were implemented using the Ax v.0.3.2 and Botorch v0.8.4 library in Python [75], and default parameters were used. Several additional scripts used in intermediate steps for formatting data tables and are described in the supplementary material. Surface plots, principal component analysis (PCA) and radar charts to explore and analyse the data were implemented using the matplotlib, seaborn and plotly packages in Python. PCA biplot was generated using the pca package in Python [76]. The surfactin molecule diagram was generated using the Pikachu package [77].

## Acknowledgements

RVA was supported by a PhD studentship from the Darwin Trust of Edinburgh. DAO was supported by the United Kingdom Research and Innovation (grant EP/S02431X/1). KB was supported by the Engineering and Physical Sciences Research Council (grant EP/V042882/1). Access to the mass spectrometry equipment was provided by EdinOmics.

## Data availability

The raw data from the QqQ-MS runs, code scripts, microplate experiment layouts, and intermediate data tables are deposited in Zenodo. Additionally, the scripts are available at the Github repository.

## Author contributions

RVA: Conceptualization, Methodology, Software, Validation, Formal analysis, Investigation, Visualisation, Data Curation, Writing – Original Draft, Writing – Review and Editing. DAO: Conceptualization, Methodology, Writing – Original Draft, Writing – Review and Editing, Supervision. KB: Conceptualization, Methodology, Writing – Original Draft, Writing – Review and Editing, Supervision, Resources, Funding acquisition.

## Notes

### Competing Interest Statement

The authors have declared no competing interest.

