## Supplementary Information for "Optimisation of surfactin yield in *Bacillus* using active learning and high-throughput mass spectrometry"

**Supplementary Figures**

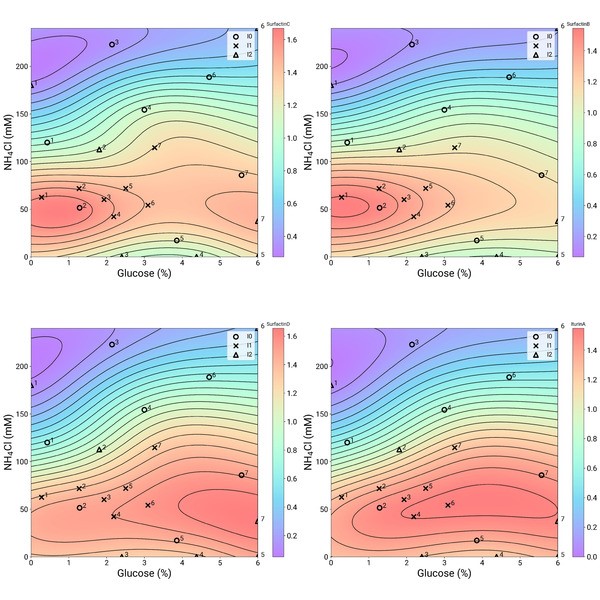

**Figure S1. Surfaces for the 4 lipopeptides that were simultaneously measures using the flow injection MS method.** Samples from all iterations are depicted in the surfaces. Colour indicates predicted titre by the Gaussian process regression model.

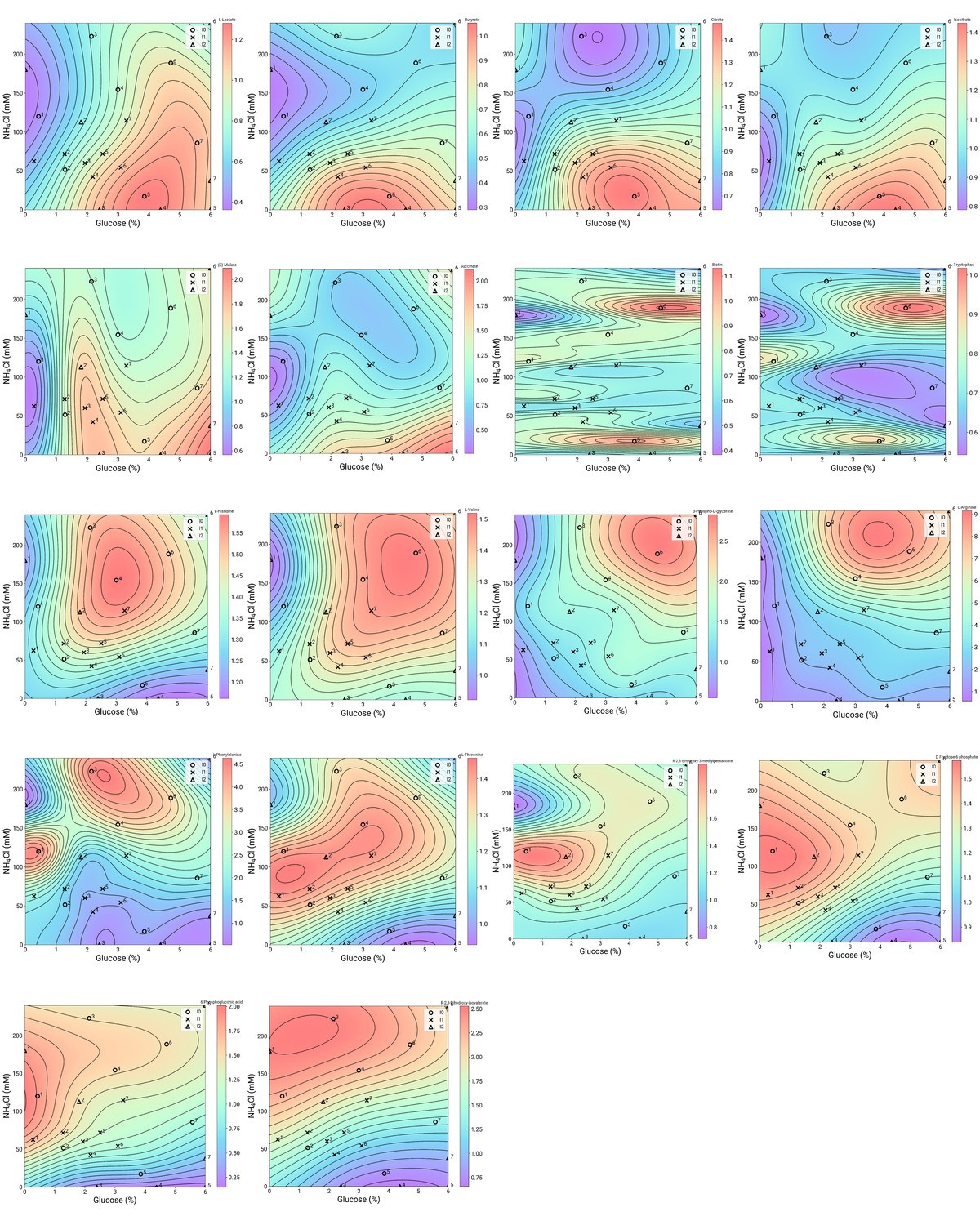

**Figure S2. Surfaces for 18 carbon/TCA-related compounds that were measured outloop using the flow injection MS method.** Samples from all iterations are depicted in the surfaces. Colour indicates predicted titre by the Gaussian process regression model.

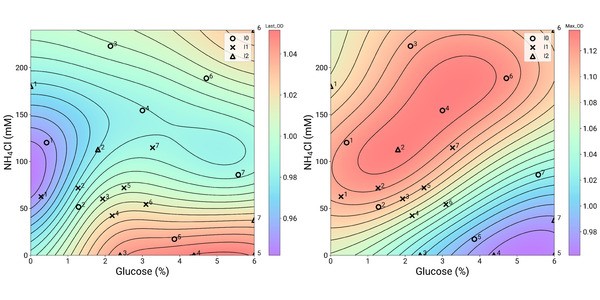

**Figure S3. Surfaces for 2 growth measurements that were obtained from the plate reader experiment.** Samples from all iterations are depicted in the surfaces. Colour indicates predicted titre by the Gaussian process regression model.

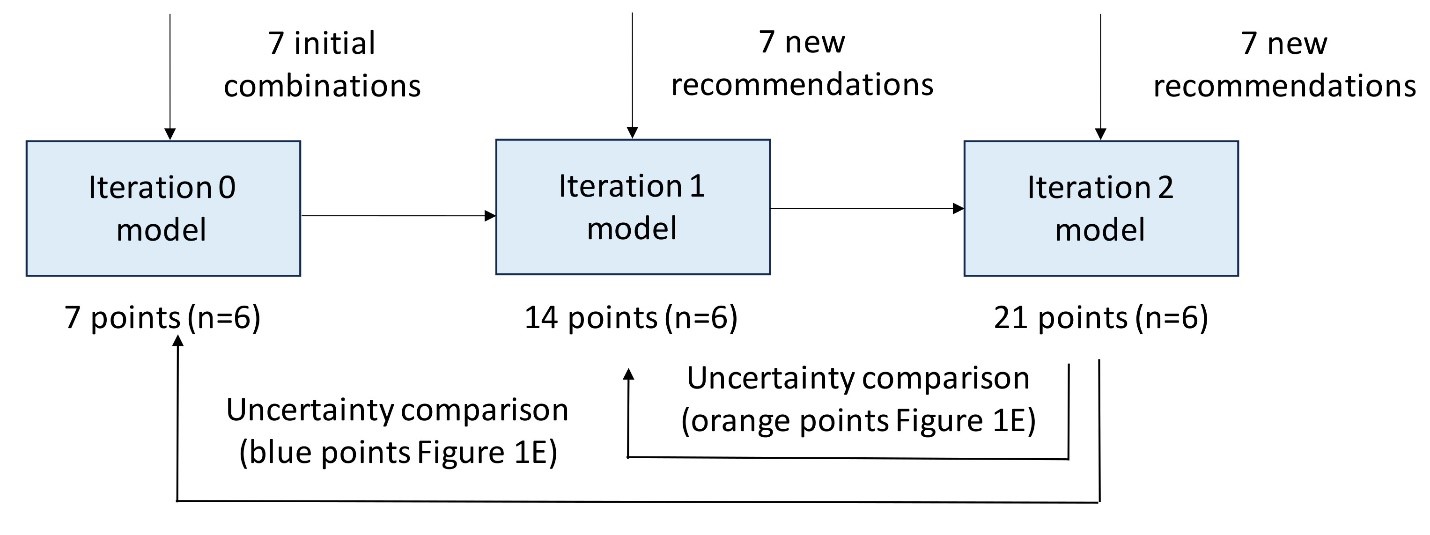

**Figure S4. Diagram showing what data serves as input to the model in each iteration and how the predicted uncertainty in the model is compared.** The model is being updated after each iteration, augmenting the available information and therefore reducing the predicted uncertainty for simulated points in the models.

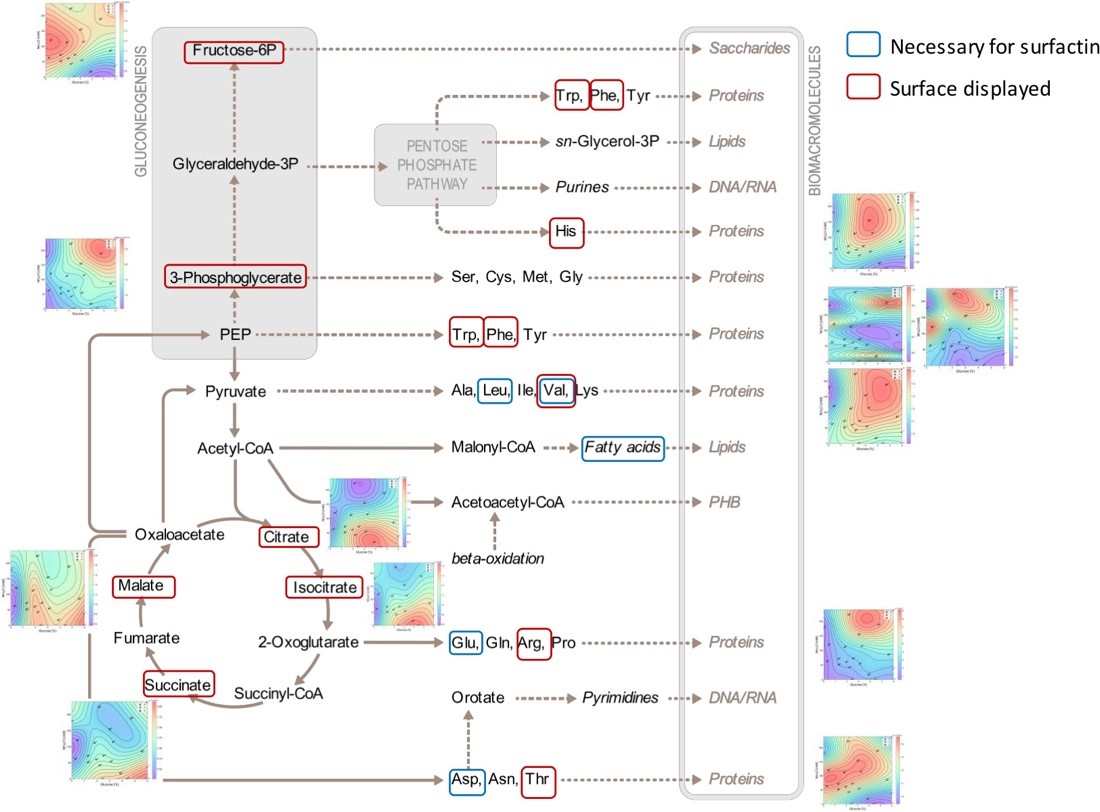

**Figure S5. Simplified pathway diagram of anabolism in *B. subtilis***, **with production surfaces assigned to the measured metabolites.** The diagram was extracted from MetaboMaps (Koblitz et al., 2020). Metabolites with available surface are enclosed with a blue box, while metabolites involved in further surfactin production are enclosed with a red box.

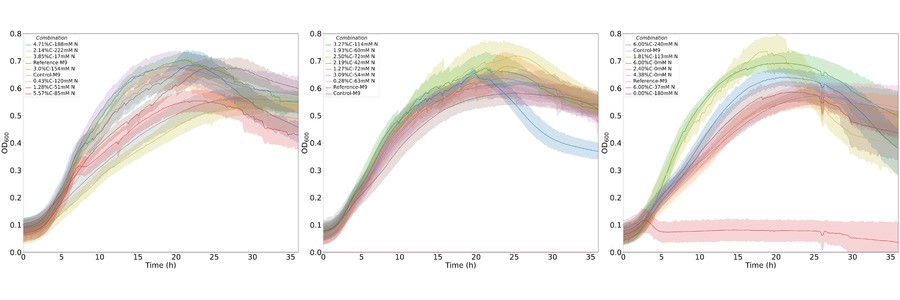

**Figure S6. Growth curves for the microplates.** From left to right, it is depicted the growth curves for different C/N combinations in Iteration 0, Iteration 1 and Iteration 2. The trend colour is given by a specific carbon and nitrogen concentration, as shown in the legend. The shadow in each trend corresponds to the confidence band calculated from the 6 biological replicates.

**Supplementary Tables**

**Table S1. Parameters used on the triple quadrupole mass spectrometer (QqQ-MS) runs using the developed flow injection method.**

| **Parameter** | **Value** |
| --- | --- |
| **Spray Voltage** | **Static** |
| **Positive Ion (V)** | **3500** |
| **Negative Ion (V)** | **3500** |
| **Sheath Gas (Arb)** | **35** |
| **Aux Gas (Arb)** | **5** |
| **Sweep Gas (Arb)** | **0** |
| **Ion transfer Tube Temp (°C)** | **325** |
| **Vaporizer Temperature (°C)** | **275** |

**Table S2. Precursor and product masses used for selected reaction monitoring (SRM) in the QqQ-MS**

| **Molecule** | **Polarity** | **Precursor mass (m/z)** | **Product mass (m/z)** | **Collision Energy (V)** | **Dwell time (ms)** |
| --- | --- | --- | --- | --- | --- |
| **Lipopeptides** | | | | | |
| SurfactinB | Positive | 1008.2 | 685 | 20 |  |
| SurfactinC | Positive | 1022.3 | 685 | 20 |  |
| IturinA | Positive | 1044.3 | 391 | 35 |  |
| SurfactinD | Positive | 1058.2 | 685 | 20 |  |
| **Central metabolism** | | | | | |
| L-Valine | Positive | 118.086 | 57.054 | 29.94 | 6.581 |
| L-Valine | Positive | 118.086 | 72 | 11.49 | 6.581 |
| L-Threonine | Positive | 120.066 | 74 | 11.4 | 6.581 |
| L-Threonine | Positive | 120.066 | 103 | 18.44 | 6.581 |
| L-Histidine | Positive | 156.077 | 93 | 23.62 | 6.581 |
| L-Histidine | Positive | 156.077 | 110.054 | 15.19 | 6.581 |
| L-Phenylalanine | Positive | 166.086 | 77 | 39.84 | 6.581 |
| L-Phenylalanine | Positive | 166.086 | 120.054 | 14.06 | 6.581 |
| L-Arginine | Positive | 175.119 | 70.071 | 23.7 | 6.581 |
| L-Arginine | Positive | 175.119 | 158.054 | 12.71 | 6.581 |
| L-Tryptophan | Positive | 205.097 | 146.071 | 18.18 | 6.581 |
| L-Tryptophan | Positive | 205.097 | 188.071 | 10.35 | 6.581 |
| L-Lactate | Negative | 89.024 | 45.125 | 12.2 | 6.581 |
| L-Lactate | Negative | 89.024 | 71.012 | 9.21 | 6.581 |
| Butyrate | Negative | 89.1 | 43.1 | 14 | 6.581 |
| Succinate | Negative | 117.019 | 73.113 | 11.74 | 6.581 |
| Succinate | Negative | 117.019 | 98.827 | 7.86 | 6.581 |
| (S)-Malate | Negative | 133.014 | 70.988 | 13.34 | 6.581 |
| (S)-Malate | Negative | 133.014 | 114.929 | 11.15 | 6.581 |
| R-2,3-Dihydroxy-isovalerate | Negative | 133.051 | 133.051 | 0 | 6.581 |
| R-2,3-dihydroxy-3-methylpentanoate | Negative | 147.066 | 147.066 | 0 | 6.581 |
| 3-Phospho-D-glycerate | Negative | 184.986 | 123.107 | 17.01 | 6.581 |
| 3-Phospho-D-glycerate | Negative | 184.986 | 166.917 | 9.04 | 6.581 |
| Citrate | Negative | 191.02 | 86.845 | 17.89 | 6.581 |
| Isocitrate | Negative | 191.02 | 116.958 | 14.81 | 6.581 |
| Isocitrate | Negative | 191.02 | 172.929 | 7.86 | 6.581 |
| Biotin | Negative | 245.1 | 227 | 14 | 6.581 |
| D-Fructose 6-phosphate | Negative | 259.022 | 138.929 | 16.5 | 6.581 |
| D-Fructose 6-phosphate | Negative | 259.022 | 168.708 | 8.71 | 6.581 |
| 6-Phosphogluconic acid | Negative | 275.017 | 195.143 | 21.68 | 6.581 |
| 6-Phosphogluconic acid | Negative | 275.017 | 256.899 | 12.83 | 6.581 |
